# Novel Anti-virulence Compounds Disrupt Exotoxin Expression in MRSA

**DOI:** 10.1101/2024.05.15.594412

**Authors:** Halie Balogh, Amaiya Anthony, Robin Stempel, Lauren Vossen, Victoria A. Federico, Gabriel Z. Valenzano, Meghan S. Blackledge, Heather B. Miller

## Abstract

Hemolysins are lytic exotoxins expressed in most strains of *S. aureus*, but hemolytic activity varies between strains. We have previously reported several novel anti-virulence compounds that disrupt the *S. aureus* transcriptome, including hemolysin gene expression. This report delves further into our two lead compounds, loratadine and a structurally related brominated carbazole, and their effects on hemolysin production in MRSA. To gain understanding into how these compounds affect hemolysis, we analyzed these exotoxins at the DNA, RNA, and protein level after in vitro treatment. While lysis of red blood cells varied between strains, DNA sequence variation did not account for it. We hypothesized that our compounds would modulate gene expression of multiple hemolysins in a laboratory strain and a clinically relevant hospital-acquired strain of MRSA, both with SCC*mec* type II. RNA-seq analysis of differential gene expression in untreated and compound-treated cultures revealed hundreds of differentially expressed genes, with a significant enrichment in genes involved in hemolysis. The brominated carbazole and loratadine both displayed the ability to reduce hemolysis in the laboratory strain, but displayed differential activity in a hospital-acquired strain. These results corroborate gene expression studies as well as western blots of alpha hemolysin. Together, this work suggests that small molecules may alter exotoxin production in MRSA, but that the directionality and/or magnitude of the difference is likely strain-dependent.

## Introduction

*Staphylococcus aureus* employs a wide range of virulence factors to establish and maintain infection in a host. The multitude of immune evasion mechanisms makes *S. aureus* infections (including those of the skin and soft tissue, bacteremia, and pneumonia) difficult to treat or prevent. Hemolysins are a group of cytolytic toxins that are expressed and secreted by bacteria to lyse host red blood cells. While only some *S. aureus* strains have acquired antibiotic resistance, almost all strains use hemolysins to enhance virulence in a host. Accordingly, hemolysins have recently become attractive targets for vaccines and anti-virulence therapies.^1–6^

*S. aureus* can express four different sets of hemolysins as virulence factors: alpha hemolysin, beta hemolysin, gamma hemolysins, and delta hemolysin. Alpha hemolysin, or alpha toxin, is the most heavily studied and well characterized. It is a pore-forming toxin that most strains encode. The gene (*hla* or *hly*) encodes a 319 amino acid polypeptide^7^ that is processed into a 293 amino acid polypeptide once a signal sequence for secretion is cleaved. The mature Hla protein has a molecular weight of approximately 33 kDa.^8^ High levels of alpha hemolysin expression have been shown to contribute to virulence in mouse models of infection with multiple strains of *S. aureus*^1^ and to the exceptional virulence of multiple community-acquired strains.^9, 10^ Alpha hemolysin also plays a direct role in the severity of infections including skin infections and sepsis.^11^

Beta hemolysin is a non pore-forming hemolysin^12^ that is not as well-studied as alpha. The *hlb* gene encodes a 330 amino acid protein^13^ (also known as phospholipase C, a sphingomyelinase)^14,15^ which gets cleaved upon secretion to become a mature protein with a molecular weight of approximately 35 kDa. Beta hemolysin is also referred to as the “hot-cold” hemolysin, as its lytic effect on red blood cells is absent or limited at 37 °C incubation, then rapidly lyses cells at 4 °C.^12^ In addition to this toxin’s sphingomyelinase activity, it also has a distinct DNA ligase biofilm activity. Hlb helps form covalent cross-links in the presence of extracellular DNA (eDNA), which strongly stimulates biofilm formation in a rabbit model of infectious endocarditis^16^ and promotes skin colonization.^17^ Furthermore, *hlb* is often interrupted by integration of bacteriophage sequence in *S. aureus*, preventing its expression unless excised.^17–19^

Gamma hemolysins are bi-component toxins that are encoded by three separate genes: *hlgA*, *hlgB*, and *hlgC.* The *hlgA* gene is transcribed separately, whereas *hlgC* and *hlgB* are transcribed as part of the same operon. These are pore-forming toxins, also produced in most strains.^20^ HlgA, B, and C are 309, 325, and 315 amino acids in length, respectively.^21^

Delta hemolysin (*hld*) is encoded within the RNAIII regulatory sequence and is also considered a phenol-soluble modulin (PSM). In *S. aureus*, it is translated as a 44 amino acid precursor with a final peptide length of only 26 amino acids.^22^

Strains of *S. aureus* have long been known to demonstrate varying hemolytic activity both in vitro and in vivo. Even *S. aureus* isolates within the same clonal lineage can vary in their degree of hemolysis. This suggests that strain-specific regulation of hemolysins exists. Whether the hemolytic activity correlates with virulence is a more complicated issue, for several reasons.^23^ First, encoding a hemolysin does not guarantee expression or the level to which it is expressed as mRNA and/or protein. Second, multiple hemolysins are produced that collectively contribute to hemolysis. These hemolysins can even act synergistically to lyse host red blood cells, as is the case with beta and delta hemolysin.^24^ Finally, hemolysins are just one facet of virulence, so other factors including exotoxins like leukocidins (including Panton-Valentine leukocidin toxins), toxic shock syndrome toxin, and phenol-soluble modulins also contribute to the arsenal of virulence factors possible in *S. aureus*.

In previous reports by our group, we demonstrated putative serine-threonine kinase (Stk1) inhibitors modulating many virulence factors in MRSA, including hemolysin genes. Our lead compound (compound 8) is a brominated carbazole that showed potentiation of multiple β-lactam antibiotics including oxacillin against several different strains of MRSA in vitro.^25^ The structurally related tricyclic antihistamine loratadine was shown to be an even more potent antibiotic adjuvant, lowering minimum inhibitory concentrations of oxacillin 32-to 512-fold.^26^ In addition to antibiotic potentiation, loratadine alone minimized biofilm formation ^26, 27^ and enhanced animal survival in a *C. elegans* infection model with multiple strains of MRSA.^27^ The transcriptome-wide effects of these compounds were measured in lab strain 43300, showing hundreds of differentially expressed genes (DEGs) compared to untreated cultures. Notably, loratadine treatment alone resulted in the modulation of multiple hemolysin genes, including *hla* (alpha hemolysin), *hlb* (beta hemolysin), *hlgA*, *hlgB*, and *hlgC* (gamma hemolysins),^28^ while compound 8 modulated *hla* levels only.^29^

Therefore, we examined with higher resolution the contributions of individual hemolysin genes in the varying hemolytic activities across strains of MRSA. We hypothesized that these compounds would result in modulation to hemolytic activity in both strains that partly contributes to overall virulence of the microorganism. These effects are in addition to antibiotic potentiation. These examinations at the DNA, RNA, and protein level would also help reveal the molecular underpinnings responsible for these strain-specific differences. In turn, these results would help guide further study, derivatization, and utility of these anti-virulence agents in MRSA.

## Results

### Hemolytic activity varies between strains of MRSA

As a qualitative examination of hemolytic activity, we evaluated the ability of several strains of MRSA to lyse red blood cells in agar petri dishes. The chosen strains represent a variety of clonal complexes. They included the laboratory reference strain, ATCC 43300, the most common hospital acquired strain in our country, USA100, a hypervirulent community acquired strain, USA300, and a clinical MRSA isolate, COL. RN4220, a methicillin sensitive strain of *S. aureus* was also included as a control, as it expresses beta, but not alpha or delta hemolysin.^30^ Importantly, alpha hemolysin is highly active against rabbit blood cells, while beta hemolysin is not.^14, 31^ Gamma hemolysin also has some activity against rabbit blood.^32^ Since gamma hemolysins are inhibited by agar,^33, 34^ the rabbit blood agar experiment primarily reports on alpha hemolysin activity. As shown in Fig. 1A, the zones of hemolysis varied in size on the rabbit blood agar plate. We concluded that USA300 showed the most alpha hemolysin activity (expression and/or secretion), followed by USA100. Strain 43300 and COL demonstrated negligible alpha hemolysin activity. As expected, RN4220 only showed a small degree of incomplete rabbit red blood cell lysis.

**Fig. 1:**
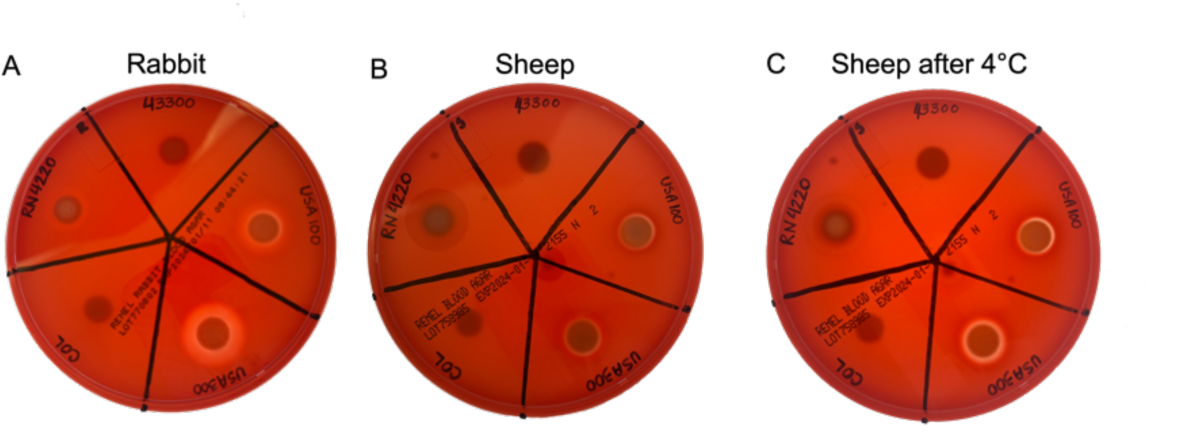
Hemolysis varies between examined strains of MRSA. (A) Each strain was grown on rabbit blood agar. (B) Each strain was grown on sheep blood agar. (C) The same sheep blood agar plate as in (B) was photographed again after 4 °C incubation.

We next examined hemolysis of sheep blood. Alpha hemolysin has limited ability to lyse sheep blood cells^35^, while beta hemolysin has high ability,^12^ so sheep blood agar primarily reports on beta hemolysin activity. As shown in Fig. 1B and 1C, USA100 and USA300 showed similar levels of beta hemolysin activity, indicated by partial lysis of cells or “bruising” that is then clear lysis after incubation at 4 °C. The 43300 and COL strains showed undetectable amounts of beta hemolysin activity. As expected, the RN4220 control showed a large hemolysis zone. Together, this indicates that the MRSA strains we are investigating differ from each other not only in hemolytic activity, but in the contribution of individual hemolysins to this activity. Using this blood agar technique, strains 43300 and COL displayed negligible alpha and beta activity, while USA100 and USA300 displayed both alpha and beta activity.

### Hemolysin genes show a high degree of sequence similarity across analyzed strains of MRSA

Sequence variation in *S. aureus* hemolysin genes exists between strains.^36, 37^ As alpha hemolysin is prominent in infection and is expressed in most *S. aureus* strains, its sequence has been analyzed in several previous publications. It was reported that *hla* is highly conserved when the gene was PCR amplified and sequenced from clinical isolates.^36, 38, 39^ To assess the level of sequence variation among the specific MRSA strains described above and utilized in our lab, we used whole-genome sequencing to take an in-depth look not only at *hla,* but all hemolysin genes. To investigate any sequence variation that would result in amino acid changes, we translated these results to construct protein alignments. As shown in Fig. S1, Hla is highly conserved, with the percent identity between pairs of strains ranging from 98.44-100%.

The gene encoding beta hemolysin (*hlb*) is often interrupted by a prophage that encodes virulence factors. Most human isolates of MRSA contain the prophage, so they do not express beta hemolysin.^18, 40^ Using whole genome sequencing, we found evidence of a prophage in *hlb* in strains 43300, USA100, and USA300. In contrast, sequencing results from COL were consistent with an uninterrupted gene. This was based on sequencing reads mapping to two locations annotated as *hlb* and analysis with PHASTEST (PHAge Search Tool with Enhanced Sequence Translation).^41^ Regardless of these strains’ ability to express the toxin, we analyzed Hlb sequences and found the percent identity between pairs of strains ranging from 95.52-100% for the upstream portion of Hlb (Fig. S2 A) and 99.27-99.64% for the downstream portion (Fig. S2 B).

There are three gamma hemolysin components, two of which showed high conservation in sequence. HlgA showed percent identity between pairs of strains ranging from 67.41-100% with COL’s sequence being most distinct from the other 3 (Fig. S3). HlgB sequences were 97.55-100% identical (Fig. S4), and HlgC sequences were 97.47-99.68% (Fig. S5).

Finally, while sequencing reads for *hld* were not annotated in our WGS results, the reference genome used for each strain showed 100% identical amino acid sequences in all pairwise comparisons (Fig. S6). Together, this indicates that the wide variation in hemolytic activity between the analyzed MRSA strains is not likely explained by DNA or protein sequence variation.

### Multiple hemolysin genes are modulated by novel anti-virulence compounds

Hemolysin gene expression varies between strains of *S. aureus*.^42–44^ The differential expression of alpha hemolysin has been particularly well-studied.^42, 43, 45, 46^ We previously reported on the putative Stk1 inhibitors compound 8 and loratadine and their in vitro effects on *S. aureus* DEGs.^27, 29^ Cultures of strain 43300 showed *hla* mRNA levels were significantly downregulated with compound 8 treatment compared to untreated controls. Other hemolysin mRNA levels did not change in a statistically significant fashion (Fig. 2).^29^ Loratadine also downregulated *hla* (Fig. 2A), while this drug upregulated *hlb* (Fig. 2B), *hlgA* (Fig. 2D), *hlgB* (Fig. 2E), and *hlgC* (Fig. 2F). These upregulated changes were large in magnitude. For example, the gamma hemolysins each showed approximately 100-fold upregulation compared to the untreated control (Fig. 2D-F). The levels of *hld* were also reduced with loratadine, but not significantly (Fig. 2C).

**Fig. 2:**
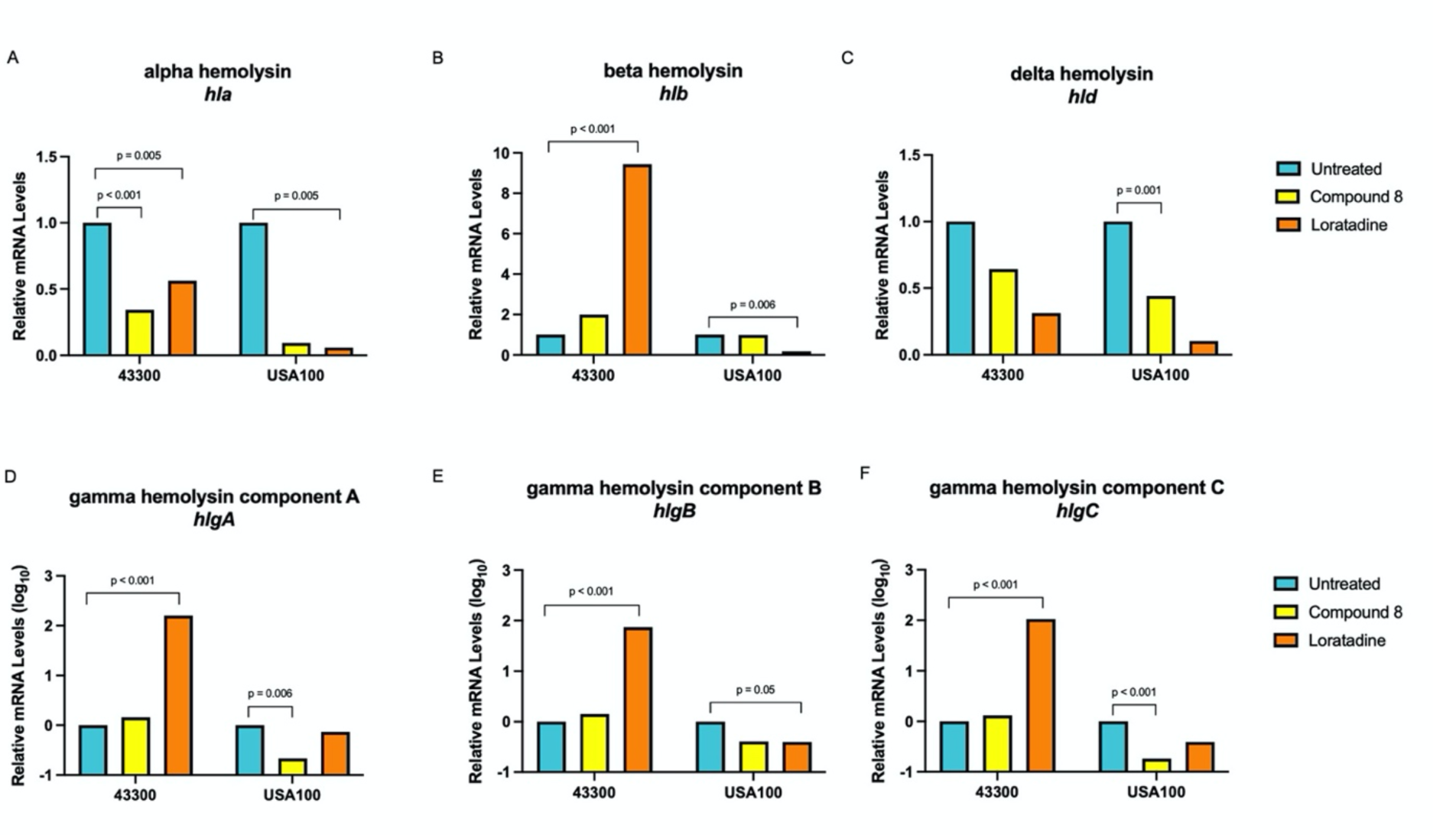
Both compound 8 and loratadine modulate gene expression of multiple hemolysins. In all panels, RNA-seq was used to determine fold changes of treated samples compared to an untreated control, set to 1. Results are averages from three independent samples. Adjusted p values ≤ 0.05 are indicated. (A) Alpha hemolysin, *hla* (B) beta hemolysin, *hlb* (C) delta hemolysin, *hld* (D) gamma hemolysin component A, *hlgA* (E) gamma hemolysin component B, *hlgB* (F) gamma hemolysin component C, *hlgC*.

We have now extended the transcriptome-wide analysis to hospital acquired USA100. Both 43300 and USA100 strains are classified as staphylococcal cassette chromosome *mec* (SCC*mec*) type II. We aimed to determine if compound 8 and/or loratadine also disrupt hemolysin gene expression in a more clinically relevant strain. Cultures of USA100 were treated in an identical fashion to what was previously reported for 43300.^27, 29^ RNA-seq sample and sequencing details are found in Tables S1-S3. This experiment revealed modulation of multiple hemolysin genes after one hour of treatment with these anti-virulence compounds. In USA100’s response to compound 8 treatment, *hla*, *hlgA, hlgB*, *hlgC*, and *hld* were downregulated. In response to loratadine, every hemolysin gene was downregulated, although not all gene expression changes were statistically significant (Fig. 2, Table S4). This repression was in contrast to laboratory reference strain 43300. Due to the fact that RNA-seq reads were present and mapped to *hlb* (shown to be interrupted by a prophage in both strains) means that the prophage was likely excised, allowing transcription. Together, this clearly indicates that these anti-virulence compounds are triggering gene expression changes in multiple hemolysins, and they are doing so in a strain-specific manner. Additionally, we conclude that loratadine generally has more pronounced effects on hemolysin gene expression than compound 8.

Due to the widespread gene expression changes in hemolysins, USA100 Gene Ontology (GO) category enrichment analysis showed that upon compound 8 treatment, DEGs belonging to the biological process of cytolysis in other organisms (GO:0051715) was significantly enriched (padj=0.006). Likewise, KEGG pathway analysis showed *Staphylococcus aureus* infection (sau05150) was significantly enriched (padj=0.006). Similar results were observed with loratadine treatment, where the *Staphylococcus aureus* infection KEGG pathway was also enriched (padj=0.02). This indicates that these anti-virulence compounds are disrupting cytotoxins more than would be expected due to chance alone.

Several of the hemolysin DEGs were validated using independently treated samples by RT-qPCR analysis (Fig. S7, Table S5). Six out of 12 tested DEG events (up or downregulation with treatment) matched between RNA-seq and RT-qPCR for a validation rate of 50%. The observation that these DEGs generally follow the same trends that were observed with the more sensitive RNA-seq method lends support that the high throughput data represents real biological changes in *S. aureus*.

Since hemolysins are primarily expressed and secreted during the post-exponential phase of growth,^47^ but our gene expression results were captured earlier during the exponential phase of growth, we repeated RT-qPCR experiments in each strain after 24 hours of growth in the presence of these compounds. After this extended treatment, repressed levels of *hla* were no longer detected with loratadine treatment in both strains tested (Fig. S3 C), illustrating a rebound to levels even higher than the untreated cultures. In contrast, the increased gene expression of *hlgA* with compound 8 and loratadine was still evident in 43300. Finally, increased gene expression of *hlgC* with compound 8 was still observed in 43300. The gamma hemolysin levels were so dramatically upregulated with just one hour of treatment that it is not surprising that levels were still higher than untreated controls after 24 hours. Natural modulation of mRNA levels due to the combination of both transcription and degradation is expected, and these data indicate that capturing mRNA levels after one hour of treatment was sufficient for detecting early gene expression changes elicited by compound 8 and loratadine. These transcript-level experiments further support the fact that these two anti-virulence compounds elicit gene expression changes in multiple hemolysins. Again, these events are not identical between the strains of MRSA tested.

### Multiple phenol-soluble modulin toxin genes are downregulated by loratadine treatment

Gamma and delta hemolysins investigated here are themselves classified as leukocidins as well, so we hypothesized that the anti-virulence compounds being studied may be affecting additional leukocidins, especially in the more virulent USA100. Bi-component leukocidin toxins (*lukDEGH*) showed no differential gene expression in USA100. Genes for *lukD* and *lukE* were not found in strain 43300, whereas genes with sequence similarity to *lukG* and *lukH* were found. Regardless, these genes did not show differential gene expression with either compound 8 nor loratadine. Additionally, leukocidins called phenol-soluble modulins (PSMs) from some strains of MRSA can lyse both red and white blood cells of the host, contributing to pathogenesis. The only PSM detected as expressed in strain 43300 was *psmβ1*, and it did not show any evidence of differential gene expression with either compound in this strain. These are more highly expressed in community then hospital acquired strains.^48^ In contrast, multiple PSMs were downregulated in the hospital acquired USA100 strain. With loratadine treatment, these cultures had reduced expression of *psmα1*, *α2*, *α3*, and *psmβ1* and *β2* compared to untreated cultures. Furthermore, loratadine lowered the expression of all four subunits of the phenol-soluble modulin transporter (*pmt*), with the downregulation of *pmtD* being statistically significant. *psm-mec*, found in the SCCmec element, did not show altered expression. None of the PSM genes or Pmt genes showed significant downregulation with compound 8 in USA100 (Table S4). This indicates that in addition to widely disrupting hemolysin gene expression, loratadine changes PSM gene expression in this hospital acquired strain.

One additional exotoxin that was examined was toxic shock syndrome toxin (TSST-1 encoded by the gene *tst*). Strain USA100 showed no evidence of expressing this toxin in any sample by RNA-seq analysis, even though this strain’s genome encodes *tst* (Table S4). However, strain 43300 showed significant downregulation of *tst* with either compound 8 or loratadine treatment compared to untreated cultures. Compound 8 reduced *tst* with a log_2_ fold change of −2.66 (padj = 6.18 x 10^-6^).^29^ Loratadine reduced *tst* even further with a log_2_ fold change of −2.98 (padj = 2.20 x 10^-30^).^27^ These results extend the number of known toxins that these compounds disrupt in the transcriptome, which could collectively lead to decreased virulence.

### In vitro hemolysis is modulated by novel anti-virulence compounds

Given the widespread disruption to multiple hemolysins’ gene expression, we next examined hemolysis levels quantitatively. MRSA strains were incubated for 24 hours with and without our lead compounds at concentrations that modulated hemolysin gene expression in our RNA-seq experiments.^27, 29^ The 24 hour incubation time allowed us to capture maximal differences in hemolysin levels as most hemolysins are produced during post-exponential growth.^46^ As shown in Fig. 3A, both untreated MRSA strains lysed rabbit red blood cells in vitro. The percent hemolysis is compared to 100% in control reactions (see Methods), indicating that the supernatants from strain 43300 showed an average of 60% hemolysis and USA100 showed an average of 52% hemolysis in these experiments.

**Fig. 3:**
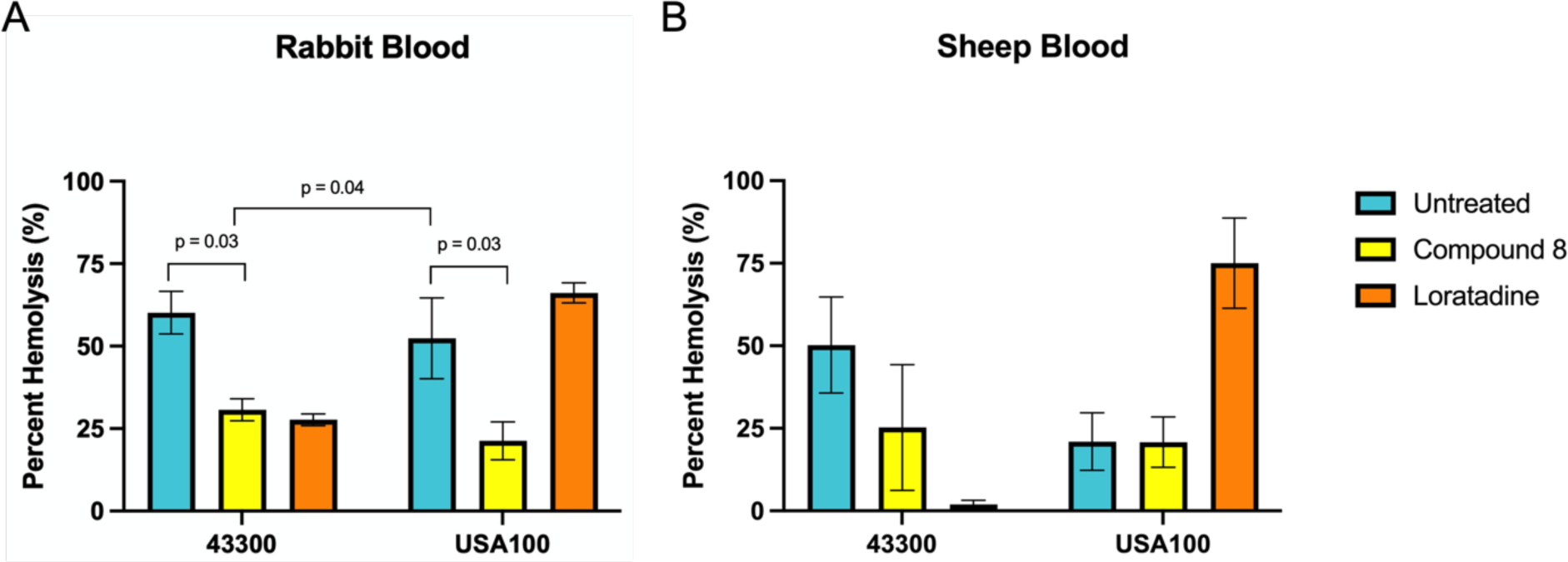
Hemolysis varies by strain and is modulated by novel anti-virulence compounds. Percent hemolysis is shown with untreated or treated samples compared to a control of blood lysed with Triton-X. Values are averages from at least three biological replicates. Error bars represent the standard error of the mean. Statistical significance was analyzed with a 2-way ANOVA comparing each mean to every other mean. Adjusted p values ≤ 0.05 are indicated. (A) Hemolysis of rabbit blood, which primarily reports alpha hemolysin activity (B) Hemolysis of sheep blood, which primarily reports beta hemolysin activity.

Compared to untreated bacterial supernatants, compound 8-treated bacterial supernatants displayed significantly reduced hemolysis due to Hla in both strains. Loratadine-treated bacterial supernatants resulted in a reduction in hemolysis in 43300, but slightly enhanced hemolysis in USA100 (66% vs 52% in the untreated control). When examining hemolysis in sheep blood (reporting beta hemolysin activity), untreated 43300 showed an average of 50% hemolysis, while USA100 showed 21% (Fig. 3B). Compared to untreated bacterial supernatants, compound 8-treated bacterial supernatants showed a decrease in sheep blood hemolysis from 50% to 25% in 43300, while USA100 remained largely unaffected. Loratadine-induced changes to hemolysis were larger in magnitude. In 43300, hemolysis decreased from 50% to 2%. USA100 showed the opposite trend, where loratadine increased hemolysis from 21% to 75%. Together, these results further support the fact that hemolysis levels differ between MRSA strains. These quantitative results also reveal what could not be measured on agar plates due to inhibitory effects of agar on gamma hemolysins.^33, 34^ Both tested strains express and secrete both alpha and beta hemolysin, but these pathogenic bacteria rely to different extents on the two toxins. In particular, USA100 demonstrated twice as much alpha hemolytic activity as beta, while 43300 produced similar amounts of each activity. In addition, loratadine seems to be a more effective modulator of both alpha and beta hemolytic activities than the smaller, 4-bromocarbazole, compound 8. USA100 showed a unique increase in both alpha and beta hemolysin activity upon loratadine treatment, although it was not statistically significant. Finally, using blood from two species which have opposite susceptibilities to alpha and beta hemolysins reveals that loratadine modulates both.

### Secreted alpha hemolysin levels are affected by novel anti-virulence compounds

Given that loratadine treatment resulted in repression of *hla* mRNA levels (Fig. 2A) that later increased (Supporting Information Fig. S7 B), and loratadine treatment increased alpha hemolysin activity in USA100 (Fig. 3A), we further analyzed Hla at the protein level. The samples used for western blotting were the same supernatants analyzed by quantitative hemolysis assays (Fig. 3). Both compound 8 and loratadine treatments resulted in slightly reduced alpha hemolysin levels in strain 43300, consistent with reduced mRNA levels (Fig. 2A) and hemolysis of rabbit blood being reduced (Fig. 3A). This reduction of Hla was not seen in USA100.

Compound 8 treated samples showed seemingly similar levels of the protein in USA100 compared to untreated samples, while loratadine treatment showed a marked increase in protein levels (Fig. 4). These western blot results are consistent with the 24 hour RT-qPCR results for *hla* (Fig. S7 C) and the rabbit blood hemolysis levels (Fig. 3A). Collectively, these results support a scenario where loratadine regulates transcription of *hla* through an unknown mechanism that results in induced expression during post-exponential phase growth when exotoxins are expressed and secreted. We observed more Hla produced by USA100 compared to 43300, consistent with blood agar plate experiments (Fig. 1A). These protein-level results also support translation and/or secretion of the toxin increasing with loratadine treatment in strain USA100, again revealing a strain-specific difference in loratadine’s effects. Finally, protein modulation of Hla is higher in magnitude with loratadine treatment compared to compound 8, consistent with our transcript-level results.

**Fig. 4:**
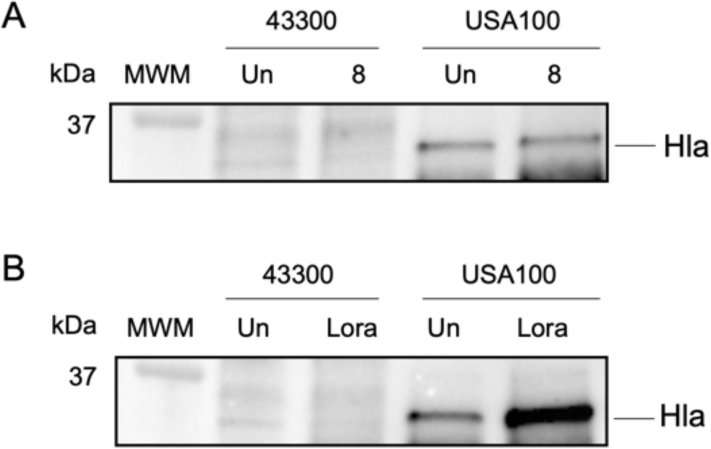
Levels of secreted alpha hemolysin are affected by 24 hours of compound 8 and loratadine treatment. In both panels, MWM, molecular weight marker; Un, untreated; 8, compound 8; Lora, loratadine. Full western blot images are found in Fig. S8.

## Discussion

Two different anti-virulence compounds, compound 8 and loratadine, are shown here to affect gene expression of multiple virulence factors in MRSA. These factors include many exotoxins including hemolysins, leukocidins, and toxic shock syndrome toxin. The regulation of these genes and proteins occurs in a strain-specific manner.

While many reports assess hemolytic activity in bacterial species, including *S. aureus*, most focus primarily on a phenotype displayed on blood agar plates. This type of analysis does not fully reveal the complex contributions that each hemolysin gene and protein product make individually. Furthermore, gamma hemolysin activities are inhibited by agar itself,^33, 34^ masking any contribution that they make in the experiment. The variation in hemolytic activity revealed by strains of interest on blood agar was consistent with a previous report.^23^ Not surprisingly, the quantitative hemolysis assays showed slightly different results than the blood agar plates. The liquid-format assays are not only more quantitative, but they do not include the inhibitory action of agar against gamma hemolysins. Therefore, while the blood agar plates quickly reveal strain-specific differences in hemolysis, the complementary quantitative hemolysis assays provide a more sensitive and accurate measurement of hemolysis. Importantly, these allow examination of anti-virulence compounds at desired concentrations in a well format that would be difficult to accurately achieve in poured agar plates.

The novel report here that loratadine downregulates *hlb* and multiple PSM genes is intriguing because these cytotoxins act synergistically to lyse host red blood cells, even in experiments using sheep blood.^49^ It may be that through this downregulation of gene expression, maximum impact on hemolysis defects (both direct and synergistic) can be imparted by loratadine.

In addition to hemolysis effects, we report here that loratadine treatment in vitro results in downregulation of several key exotoxins tied to biofilm formation: *hlb*, *psmα1*, *α*2, *α*3, and *psmβ1* and *β2*. Beta hemolysin was shown to stimulate biofilm formation in rabbit models of infection.^16, 17^ In addition, all PSMs in *S. aureus* enhance the dissemination of biofilms throughout the body. These peptides accomplish this by promoting not just structuring of biofilms, but also detachment.^50^ Accordingly, loratadine-regulated repression of these factors that promote biofilms likely helps explain in part the anti-biofilm effects we and others have previously reported.^26, 27, 51^ Additional experiments will be needed to analyze the detailed mechanism through which loratadine works against biofilms.

Each of the experiments presented here demonstrates that loratadine disrupts cytotoxin genes differently in 43300 compared to USA100. Surprisingly, loratadine increased alpha and beta hemolytic activity in USA100 but not 43300 (Fig. 3, Fig. 4). In USA100, the loratadine-regulated repression of toxin genes including PSMs may compensate for the upregulated hemolysis we are reporting. Additional investigations will focus on this strain-specific regulation, the conditions and timing in which it occurs, and whether it extends to other strains of MRSA.

While our previous reports have provided putative evidence that compound 8 and loratadine interact with and inhibit Stk1,^25–27, 29^ we have not comprehensively shown that Stk1 is the only target of these compounds. It is possible that one or both of these compounds is promiscuous and that interaction with one or more additional molecular targets in MRSA contributes to the observed effects on hemolysin and virulence gene expression. Candidate proteins are currently being examined to learn more about these anti-virulence compounds’ action in clinically relevant strains of *S. aureus*.

## Methods

### Bacterial strains

*S. aureus* 43300 and USA100 (BAA-1753) were purchased from the American Type Culture Collection (ATCC). *S. aureus* RN4220 was purchased from BEI Resources.

### Blood agar assays

MRSA strains were purchased from grown overnight in cation-adjusted Mueller-Hinton broth (CAMHB) at 37 °C for 16 hours with shaking at 220 rpm. Overnight cultures were pipetted on the surface of sheep blood agar plates and incubated at 37 °C for 18 hours and then at 4 °C for 18 hours before being photographed on a white light box. The same method was used for rabbit blood agar plates except that incubation was at 37 °C for 18 hours only.

### Whole genome sequencing

MRSA strains were grown in triplicate cultures overnight in CAMHB at 37 °C for 16 hours with shaking at 220 rpm. Genomic DNA was purified using a Monarch gDNA purification kit from New England Biolabs. Cell lysis buffer was supplemented with lysostaphin. The integrity of each sample was assessed via agarose gel electrophoresis and imaged on a ChemiDoc by BioRad. Sample concentration and A_260_/A_280_ were measured by a Nanodrop lite by Thermo Scientific. Seqcenter performed Illumina DNA library preparation and whole genome sequencing. PHASTEST (PHAge Search Tool with Enhanced Sequence Translation) was used to detect prophage sequences.^41^

### Amino acid alignments

Whole genome sequencing paired reads were imported into Kbase^52^ where they were assembled using the SPADES app and annotated using the RAST app. Genes of interest were pasted into Clustal Omega for alignment. Jalview was used to translate into amino acids and create the alignments shown. Jalview was used to calculate the percent identities.

### RNA-sequencing

MRSA cultures, treatments, and RNA purification were performed as described previously.^53^ All subsequent RNA-seq steps, including additional RNA quality control (Table S1), library construction, sequencing (Table S2), reference genome mapping (Table S3) and differential expression analysis (Table S4) were conducted by Novogene, Inc as reported previously. Results reported here in strain ATCC 43300 are from an RNA-seq data set previously described concerning compound 8^53^ and loratadine treatment.^27^ Experimental details, raw, and processed data were deposited to Gene Expression Omnibus (GEO) for strain 43300 with loratadine treatment under accession number GSE227099, and with compound 8 treatment under accession number GSE193395. Results reported here in strain USA100 with loratadine and/or compound 8 treatment are under accession number GSE267020.

### RT-qPCR

Total RNA was purified from treated bacterial cultures, reverse transcribed, and amplified with gene-specific primers and relative gene expression was determined as previously described.^26^ These samples were independent from those analyzed by RNA-seq. Primer sequences and calculated efficiencies are in Table S5.

### Quantitative hemolysis assays

Assays were based on those previously published.^54^ Cells were cultured overnight in CAMHB. Overnight cultures were diluted 1:200 in 5ml of fresh CAMHB containing the indicated compounds or antibiotic or untreated and were grown at 37 °C with shaking for 24 hours. Cultures were centrifuged and supernatants were isolated and frozen at −80 °C overnight before use. Erythrocytes were freshly diluted to 4% solutions in PBS. In a 96-well plate, 100 μL of erythrocytes were mixed with 100 μL of supernatants. Plates were incubated at 37 °C for 60 minutes and the absorbance was recorded at 550nm on a Biotek synergy H1 spectrophotometer. PBS was used as a negative (0 % hemolysis) control and 0.1% Triton-X was used as a positive (100% hemolysis) control. Technical replicates were performed in at least triplicate. Biological replicates were performed in at least triplicate.

### Western blots

Supernatants described above were also used in western blot experiments. Each supernatant was already normalized to the same OD, so equal volumes were used. After denaturing samples at 95 °C for 10 minutes, they were loaded onto an SDS-PAGE gel (BioRad any kD stain-free gels). After electrophoresis, gels were transferred to PVDF membranes using a BioRad Transblot Turbo and transfer efficiency was assessed on a BioRad ChemiDoc imager using the stain-free technology. Membranes were blocked in 5% nonfat dry milk in Tris-buffered saline with 1% Tween-20 (TBS-T) at room temperature for 30 minutes. After three washes in TBS-T, the primary antibody (rabbit anti-*Staph* alpha toxin from Sigma) was used at a 1:10,000 dilution in TBS-T, rocking overnight at 4 °C. After three additional washes in TBS-T, the secondary antibody (goat anti-rabbit Horse Radish Peroxidase from Invitrogen) was used at a 1:5,000 dilution in TBS-T, rocking for 30 minutes at room temperature. Final TBS-T washes were followed by development with BioRad Clarity ECL. Western blots were imaged with a BioRad ChemiDoc imager.

### Statistical Analyses

Statistical significance in RT-qPCR experiments and quantitative hemolysis assays was determined using one-way and two-way ANOVAs, respectively, with Tukey’s multiple comparisons test in GraphPad Prism. RT-qPCR data comparing only two treatments used an unpaired student’s t-test in GraphPad Prism.

## Note

The authors declare the following competing financial interest(s): M.S.B. and H.B.M. have filed a patent on the technology disclosed.

## Supporting information

Balogh et al Supporting Information Fig S1-S8 Tables S1-S3,S5

Balogh et al Supporting Information Table S4

## Acknowledgements

This work was funded by the National Institutes of Health (R15GM134503 to M.S.B. and H.B.M.) and High Point University.

