## Supplementary material for "Novel Anti-virulence Compounds Disrupt Exotoxin Expression in MRSA": Balogh et al Supporting Information Fig S1-S8 Tables S1-S3,S5

Page

**Supporting Tables**
S1 – Additional quality control details of USA100 RNA samples 2
S2 – RNA-seq read quality control 3
S3 – Mapped reads summary 4
S4 – All differential gene expression results for USA100 (.xlsx)
S5 – RT-qPCR primer information 5

**Supporting Figures**
S1 – Alpha hemolysin precursor alignment 6
S2 – Beta hemolysin alignment 6
S3 – Gamma hemolysin component A alignment 7
S4 - Gamma hemolysin component B alignment 7
S5 - Gamma hemolysin component C alignment 7
S6 – Delta hemolysin precursor alignment 8
S7 - RT-qPCR validation of RNA-seq determined gene expression changes 8
S8 – Full western blot images 9

References 10

**Table S1: Additional quality control details of USA100 RNA samples.** Un represents untreated sample, ox represents oxacillin treated sample, 8 represents compound 8 treated sample, lor represents loratadine treated sample, 8ox represents a cotreated sample, and lorox represents a cotreated sample. Each biological replicate is labelled A, B, or C. RIN= RNA integrity number.

| Sample # | Sample ID | Customer Sample ID | Sample Type | Sample Volume (ul) | Concentration (ng/ul) | Total Quantity (ng) | RIN |
|---|---|---|---|---|---|---|---|
| 1 | 1735R-2069-01 | USA100unA | RNA | 53 | 170.9 | 9055.9 | 7.6 |
| 2 | 1735R-2069-02 | USA100oxA | RNA | 53 | 44.3 | 2346.0 | 7.6 |
| 3 | 1735R-2069-03 | USA1008A | RNA | 52 | 57.1 | 2968.5 | 7.1 |
| 4 | 1735R-2069-04 | USA100lorA | RNA | 52 | 123.4 | 6418.8 | 7.3 |
| 5 | 1735R-2069-05 | USA1008oxA | RNA | 52 | 150.6 | 7830.2 | 7.2 |
| 6 | 1735R-2069-06 | USA100loroxA | RNA | 65 | 8.8 | 573.7 | NA |
| 7 | 1735R-2069-07 | USA100unB | RNA | 52 | 370.6 | 19271.1 | 7.9 |
| 8 | 1735R-2069-08 | USA100oxB | RNA | 52 | 28.0 | 1456.4 | 7.1 |
| 9 | 1735R-2069-09 | USA1008B | RNA | 52 | 85.6 | 4452.5 | 7.0 |
| 10 | 1735R-2069-10 | USA100lorB | RNA | 52 | 97.6 | 5076.5 | 6.8 |
| 11 | 1735R-2069-11 | USA1008oxB | RNA | 52 | 39.2 | 2040.7 | 7.5 |
| 12 | 1735R-2069-12 | USA100loroxB | RNA | 52 | 55.2 | 2869.6 | 6.7 |
| 13 | 1735R-2069-13 | USA100unC | RNA | 51 | 267.5 | 13640.8 | 8.1 |
| 14 | 1735R-2069-14 | USA100oxC | RNA | 48 | 6.7 | 322.0 | NA |
| 15 | 1735R-2069-15 | USA1008C | RNA | 51 | 41.8 | 2134.0 | 7.2 |
| 16 | 1735R-2069-16 | USA100lorC | RNA | 51 | 103.5 | 5280.3 | 7.3 |
| 17 | 1735R-2069-17 | USA1008oxC | RNA | 51 | 146.1 | 7448.8 | 7.3 |
| 18 | 1735R-2069-18 | USA100loroxB | RNA | 48 | 5.5 | 264.8 | NA |

**Table S2: RNA-seq reads quality control details.** Un represents untreated sample, ox represents oxacillin treated sample, Cmpd8 represents compound 8 treated sample, Lor represents loratadine treated sample, Cmpd8_Ox represents cotreated sample, and Lor_Ox represents cotreated sample. Each biological replicate is labelled A, B, or C. Q20 and Q30 were calculated as the base number of Phred value > 20 or 30, respectively, divided by the total base value x 100%.

| Sample name | Raw reads | Clean reads | Raw bases | Clean bases | Error rate | Q20 | Q30 | GC content |
|---|---|---|---|---|---|---|---|---|
| Un_A | 16563404 | 16199626 | 2.49G | 2.43G | 0.03 | 97.55 | 93.01 | 34.95 |
| Ox_A | 16407200 | 15973518 | 2.47G | 2.4G | 0.03 | 97.61 | 93.16 | 35.00 |
| Cmpd8_A | 17782084 | 17355948 | 2.67G | 2.61G | 0.03 | 97.57 | 93.06 | 35.38 |
| Lor_A | 17763720 | 17237008 | 2.67G | 2.59G | 0.03 | 97.63 | 93.18 | 34.99 |
| Cmpd8_Ox_A | 18616198 | 18165322 | 2.8G | 2.73G | 0.03 | 97.57 | 93.05 | 35.30 |
| Lor_Ox_A | 17264464 | 16941150 | 2.59G | 2.55G | 0.03 | 97.59 | 93.1 | 34.94 |
| Un_B | 18606480 | 18109918 | 2.8G | 2.72G | 0.03 | 97.49 | 92.87 | 35.13 |
| Ox_B | 16118996 | 15664202 | 2.42G | 2.35G | 0.03 | 97.59 | 93.1 | 34.79 |
| Cmpd8_B | 17760996 | 17308642 | 2.67G | 2.6G | 0.03 | 97.68 | 93.3 | 35.48 |
| Lor_B | 16010694 | 15431796 | 2.41G | 2.32G | 0.03 | 97.71 | 93.37 | 35.01 |
| Cmpd8_Ox_B | 14668862 | 14310466 | 2.21G | 2.15G | 0.03 | 97.69 | 93.32 | 35.25 |
| Lor_Ox_B | 13825412 | 13380550 | 2.08G | 2.01G | 0.03 | 97.8 | 93.65 | 34.85 |
| Un_C | 14488574 | 14014188 | 2.18G | 2.11G | 0.03 | 97.64 | 93.19 | 34.75 |
| Ox_C | 14465062 | 13989522 | 2.17G | 2.1G | 0.03 | 97.75 | 93.53 | 35.28 |
| Cmpd8_C | 14354250 | 13838842 | 2.16G | 2.08G | 0.03 | 97.76 | 93.5 | 34.97 |
| Lor_C | 12894550 | 12528502 | 1.94G | 1.88G | 0.03 | 97.74 | 93.45 | 34.87 |
| Cmpd8_Ox_C | 15374400 | 14948966 | 2.31G | 2.25G | 0.03 | 97.61 | 93.2 | 35.41 |
| Lor_Ox_C | 13252778 | 12708422 | 1.99G | 1.91G | 0.03 | 97.56 | 93.26 | 36.32 |

**Supporting Table S3: Mapped reads summary.** Un represents untreated sample, ox represents oxacillin treated sample, Cmpd8 represents compound 8 treated sample, Lor represents loratadine treated sample, Cmpd8_Ox represents cotreated sample, and Lor_Ox represents cotreated sample. Each biological replicate is labelled A, B, or C.

| **Sample name** | **Cmpd8_A** | **Cmpd8_B** | **Cmpd8_C** | **Cmpd8_Ox_A** | **Cmpd8_Ox_B** | **Cmpd8_Ox_C** | **Lor_A** | **Lor_B** | **Lor_C** |
|---|---|---|---|---|---|---|---|---|---|
| Total reads | 17355948 | 17308642 | 13838842 | 18165322 | 14310466 | 14948966 | 17237008 | 15431796 | 12528502 |
| Total mapped | 17124337 (98.67%) | 16745919 (96.75%) | 13632347 (98.51%) | 17917673 (98.64%) | 13821016 (96.58%) | 14726483 (98.51%) | 17004296 (98.65%) | 14877913 (96.41%) | 12354208 (98.61%) |
| Multiple mapped | 653694 (3.77%) | 916949 (5.3%) | 501832 (3.63%) | 930892 (5.12%) | 723524 (5.06%) | 837940 (5.61%) | 572647 (3.32%) | 683508 (4.43%) | 426828 (3.41%) |
| Uniquely mapped | 16470643 (94.9%) | 15828970 (91.45%) | 13130515 (94.88%) | 16986781 (93.51%) | 13097492 (91.52%) | 13888543 (92.91%) | 16431649 (95.33%) | 14194405 (91.98%) | 11927380 (95.2%) |
| Read-1 | 8240255 (47.48%) | 7922309 (45.77%) | 6567562 (47.46%) | 8498666 (46.79%) | 6554086 (45.8%) | 6947681 (46.48%) | 8220857 (47.69%) | 7102905 (46.03%) | 5966217 (47.62%) |
| Read-2 | 8230388 (47.42%) | 7906661 (45.68%) | 6562953 (47.42%) | 8488115 (46.73%) | 6543406 (45.72%) | 6940862 (46.43%) | 8210792 (47.63%) | 7091500 (45.95%) | 5961163 (47.58%) |
| Reads map to '+' | 8236092 (47.45%) | 7917616 (45.74%) | 6566198 (47.45%) | 8494025 (46.76%) | 6551387 (45.78%) | 6944313 (46.45%) | 8216744 (47.67%) | 7099985 (46.01%) | 5964315 (47.61%) |
| Reads map to '-' | 8234551 (47.45%) | 7911354 (45.71%) | 6564317 (47.43%) | 8492756 (46.75%) | 6546105 (45.74%) | 6944230 (46.45%) | 8214905 (47.66%) | 7094420 (45.97%) | 5963065 (47.6%) |
| Reads mapped in proper pairs | 15178748 (87.46%) | 14427716 (83.36%) | 12207224 (88.21%) | 15629750 (86.04%) | 12020118 (84%) | 12626678 (84.47%) | 15282116 (88.66%) | 13020262 (84.37%) | 11087146 (88.5%) |
| Proper-paired reads map to different chrom | 0 (0%) | 0 (0%) | 0 (0%) | 0 (0%) | 0 (0%) | 0 (0%) | 0 (0%) | 0 (0%) | 0 (0%) |

| **Sample name** | **Lor_Ox_A** | **Lor_Ox_B** | **Lor_Ox_C** | **Ox_A** | **Ox_B** | **Ox_C** | **Un_A** | **Un_B** | **Un_C** |
|---|---|---|---|---|---|---|---|---|---|
| Total reads | 16941150 | 13380550 | 12708422 | 15973518 | 15664202 | 13989522 | 16199626 | 18109918 | 14014188 |
| Total mapped | 16619486 (98.1%) | 12903635 (96.44%) | 12391696 (97.51%) | 15701622 (98.3%) | 15063006 (96.16%) | 13692023 (97.87%) | 15995883 (98.74%) | 17455039 (96.38%) | 13830339 (98.69%) |
| Multiple mapped | 888045 (5.24%) | 642245 (4.8%) | 1468567 (11.56%) | 634985 (3.98%) | 718424 (4.59%) | 929260 (6.64%) | 494791 (3.05%) | 1025989 (5.67%) | 412861 (2.95%) |
| Uniquely mapped | 15731441 (92.86%) | 12261390 (91.64%) | 10923129 (85.95%) | 15066637 (94.32%) | 14344582 (91.58%) | 12762763 (91.23%) | 15501092 (95.69%) | 16429050 (90.72%) | 13417478 (95.74%) |
| Read-1 | 7869562 (46.45%) | 6135011 (45.85%) | 5464181 (43%) | 7536052 (47.18%) | 7176663 (45.82%) | 6382457 (45.62%) | 7754299 (47.87%) | 8222475 (45.4%) | 6711588 (47.89%) |
| Read-2 | 7861879 (46.41%) | 6126379 (45.79%) | 5458948 (42.96%) | 7530585 (47.14%) | 7167919 (45.76%) | 6380306 (45.61%) | 7746793 (47.82%) | 8206575 (45.32%) | 6705890 (47.85%) |
| Reads map to '+' | 7867176 (46.44%) | 6133063 (45.84%) | 5461995 (42.98%) | 7535189 (47.17%) | 7173192 (45.79%) | 6382866 (45.63%) | 7751824 (47.85%) | 8216554 (45.37%) | 6709875 (47.88%) |
| Reads map to '-' | 7864265 (46.42%) | 6128327 (45.8%) | 5461134 (42.97%) | 7531448 (47.15%) | 7171390 (45.78%) | 6379897 (45.6%) | 7749268 (47.84%) | 8212496 (45.35%) | 6707603 (47.86%) |
| Reads mapped in proper pairs | 14680818 (86.66%) | 11723158 (87.61%) | 10259114 (80.73%) | 14083462 (88.17%) | 13387146 (85.46%) | 12026152 (85.97%) | 14291416 (88.22%) | 14902196 (82.29%) | 12166502 (86.82%) |
| Proper-paired reads map to different chrom | 0 (0%) | 0 (0%) | 0 (0%) | 0 (0%) | 0 (0%) | 0 (0%) | 0 (0%) | 0 (0%) | 0 (0%) |

**Supporting Information Table S5: RT-qPCR primers used in this study.**

| Gene | Forward | Reverse | Efficiency | Reference |
|---|---|---|---|---|
| *16S* | CTGTGCACATCTTGACGGTA | TCAGCGTCAGTTACAGACCA | 93.80% | Yarwood et al.[1] |
| *hla* | ATGAATCCTGTCGCTAATGCCG | TGACCAGCAATGGTACCTTTCG | 107.02% | This work |
| *hlgA* | AGCAGTTGGTTTAATCGCCCCTTT | TTGATGATTTCTGCACCTTGGCCG | 83.86% | Viering et al.[2] |
| *hlgC* | GGTGGTAATTTCCAATCAGCC | GAATGAATTCGCTTTGACGCCC | 109.18% | This work |

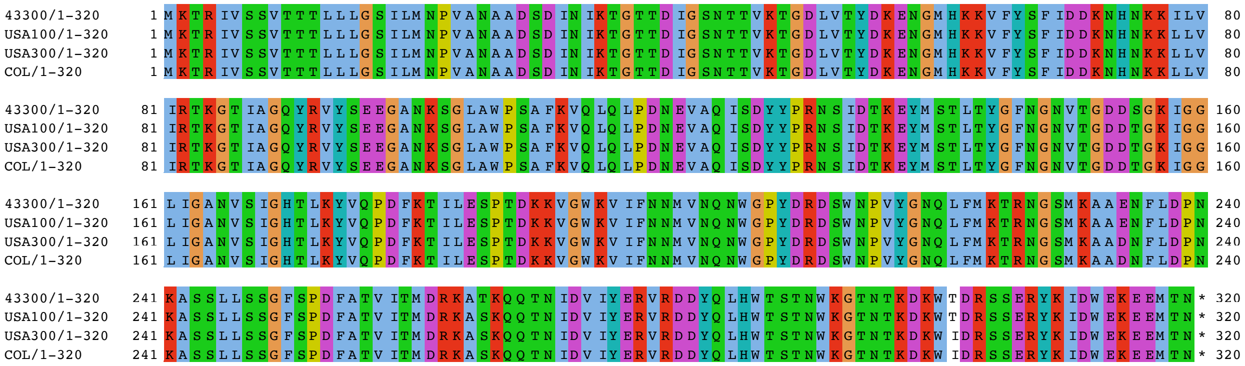


**Supporting Figure S1: Alpha hemolysin precursor is highly conserved among analyzed MRSA strains.** Amino acids are colored based on Clustal settings.

A)

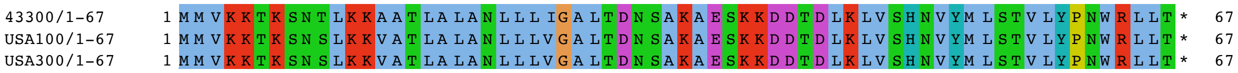


B)

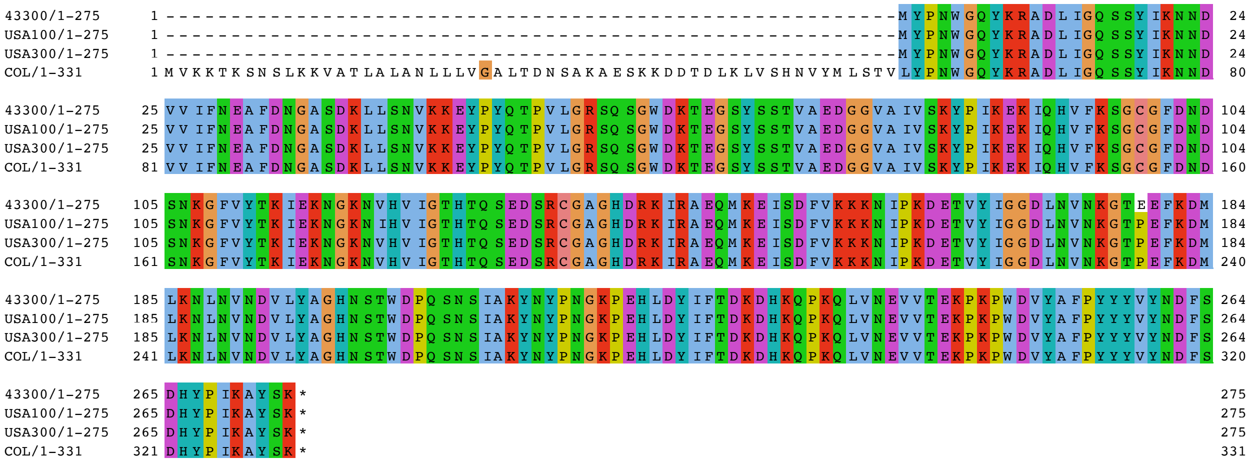


**Supporting Figure S2: Beta hemolysin is highly conserved among analyzed MRSA strains.** In both panels, amino acids are colored based on Clustal settings. A) The sequences are identical in the upstream instance (amino acids 1-67). B) This gene is disrupted by a prophage in MRSA strains 43300, USA100, and USA300, but the highly conserved downstream instance is shown.

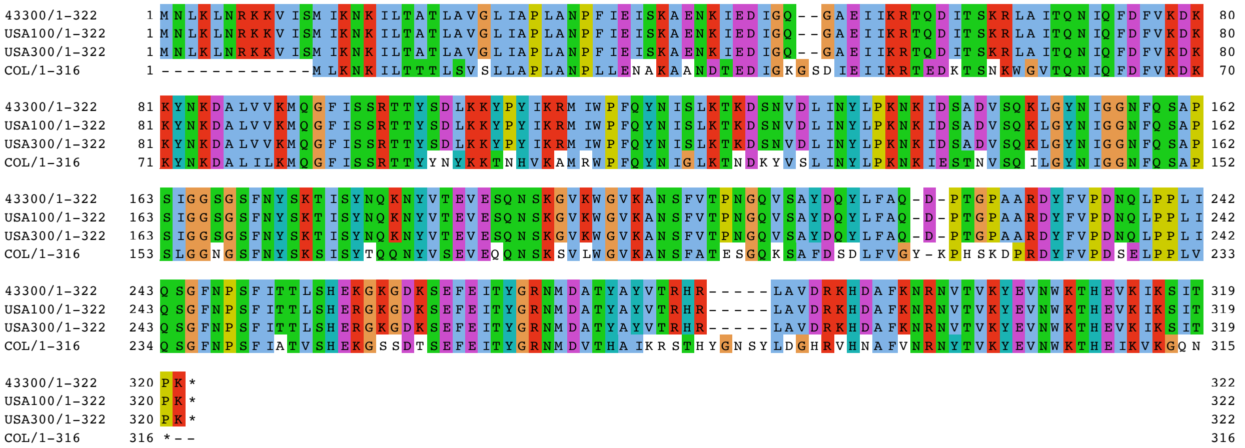

**Supporting Figure S3: Gamma hemolysin component A is highly conserved among 43300, USA100 and USA100.** Amino acids are colored based on Clustal settings.

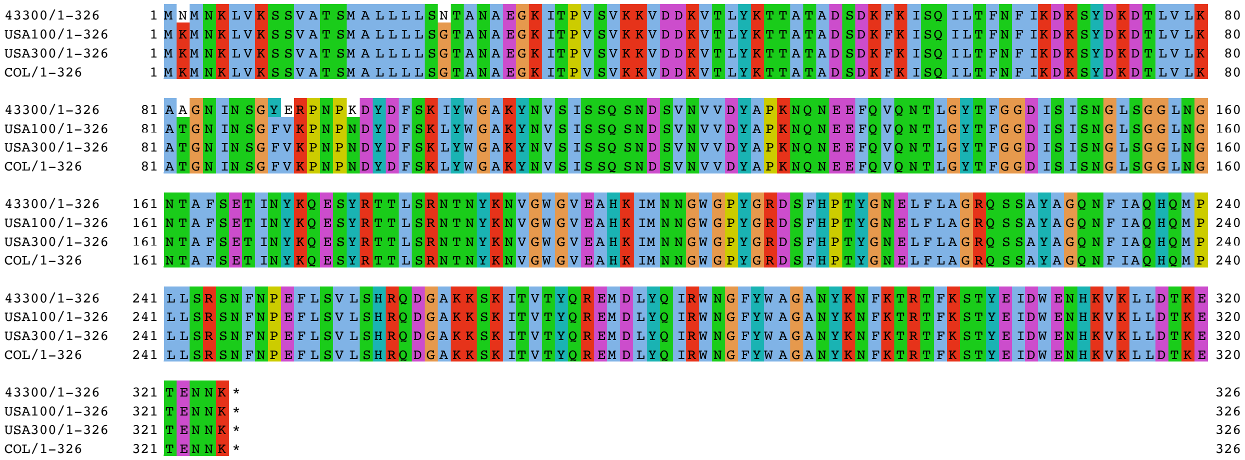

**Supporting Figure S4: Gamma hemolysin component B is highly conserved among analyzed MRSA strains.** Amino acids are colored based on Clustal settings.

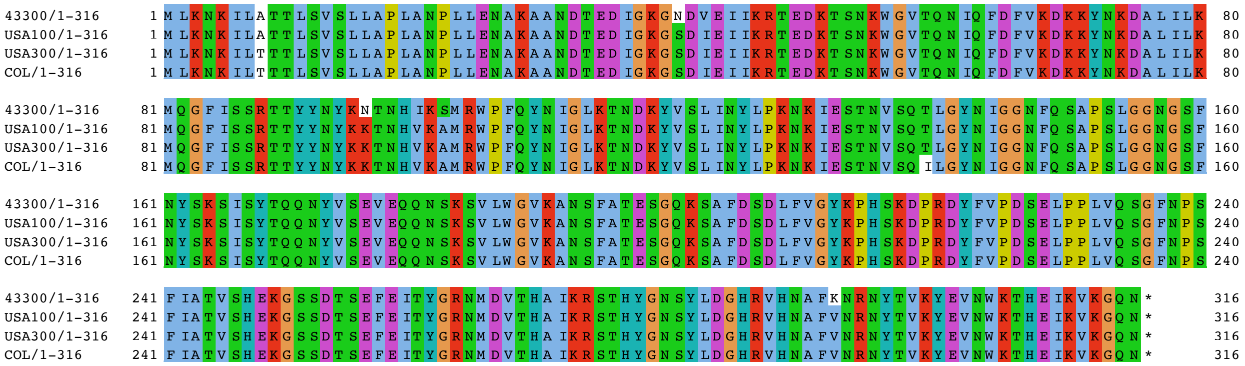

**Supporting Figure S5: Gamma hemolysin component C is highly conserved among analyzed MRSA strains.** Amino acids are colored based on Clustal settings.

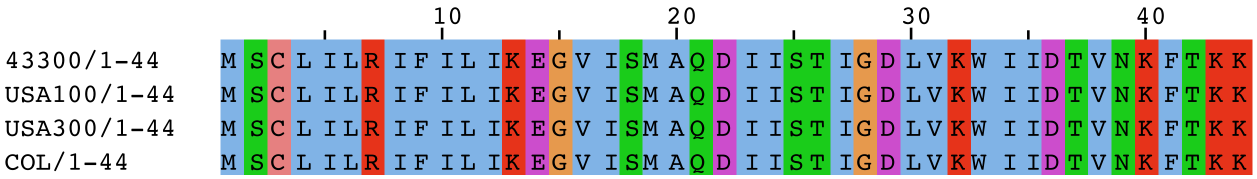


**Supporting Figure S6: Delta hemolysin precursor is identical in analyzed MRSA strains.** Amino acids are colored based on Clustal settings.

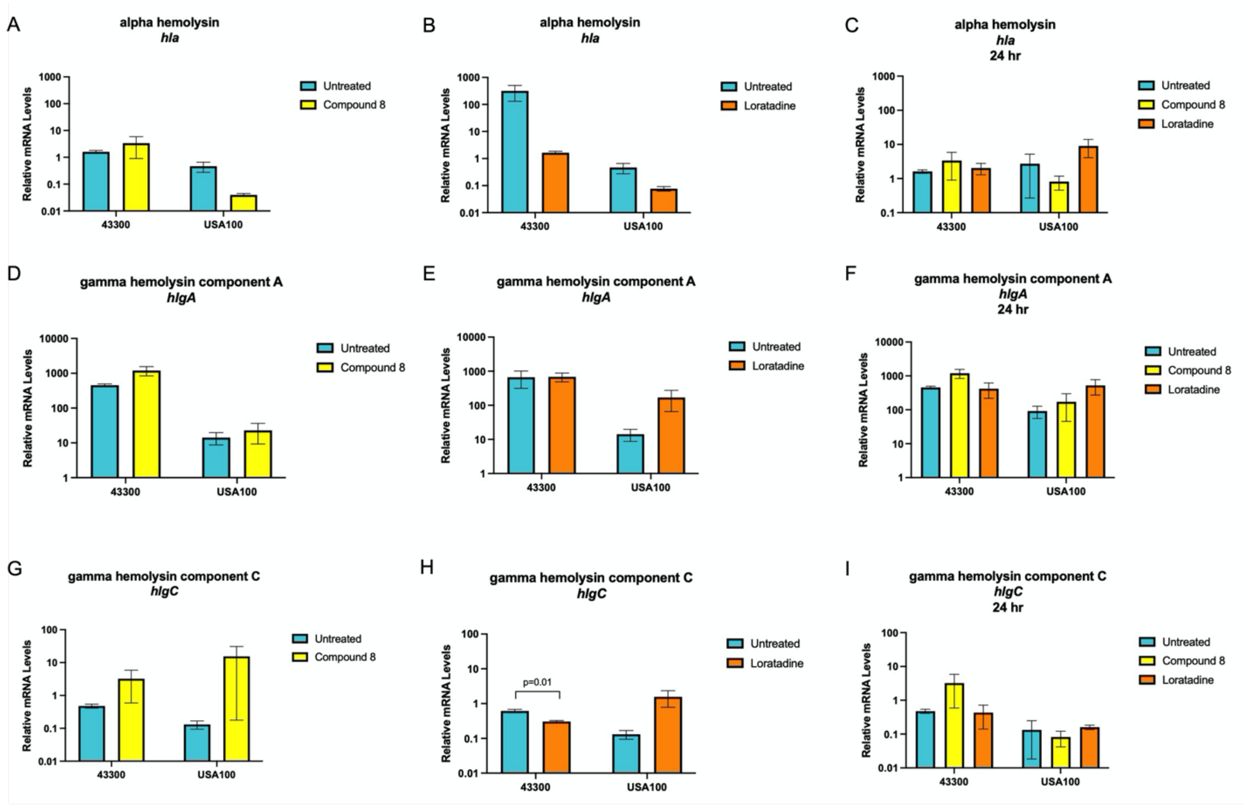


**Supporting Figure S7: RT-qPCR validation of RNA-seq determined gene expression changes.** Results are the mean of at least three biological replicates. The y axis displays the average mRNA levels relative to *16S* rRNA. Error bars represent the standard error of the mean. Statistical significance was analyzed by students unpaired t-tests between treated and untreated levels. p values ≤ 0.05 are indicated.

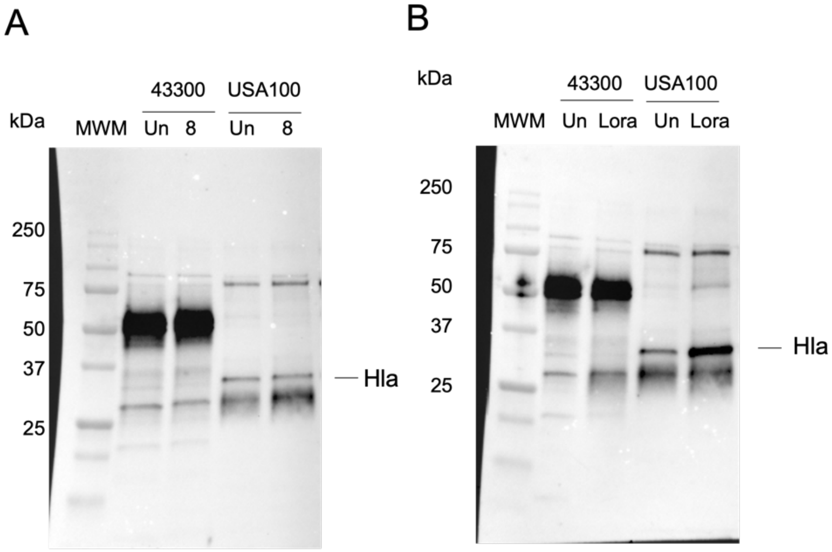


**Supporting Figure S8: Levels of secreted alpha hemolysin are affected by compound 8 and loratadine treatment.** A) Full western blot image of that shown in Figure 4A. B) Full western blot image of that shown in Figure 4B. In all panels, MWM is molecular weight marker, Un is untreated, 8 is compound 8, and Lora is loratadine.
